# Perception of audio-visual synchrony is modulated by walking speed and step-cycle phase

**DOI:** 10.1101/2024.07.21.604456

**Authors:** Gabriel Clouston, Matt Davidson, David Alais

**Author notes:** equal contribution (Gabriel Clouston, Matt Davidson).

## Abstract

Investigating sensory processes in active human observers is critical for a holistic understanding of perception. Recent research has demonstrated that locomotion can alter visual detection performance in a rhythmic manner, illustrating how a very frequent and natural behaviour can influence sensory performance. Here we extend this line of work to incorporate variations in walking speed, and test whether multi-sensory processing is impacted by the speed and phase of locomotion. Participants made audio-visual synchrony judgements while walking at two speeds over a range of stimulus onset asynchronies (SOAs). We find that sensitivity to multi-sensory synchrony decreases at slow walking speeds and is accompanied by an increase in reaction times, compared to when walking at a natural pace. A further analysis of the shortest SOAs was conducted to test whether subjective synchrony modulated over the step cycle. This revealed that synchrony judgements were quadratically modulated with perceived synchrony being higher in the swing phase of each step and lower when both feet were grounded during stance phase. Together, these results extend an earlier report that walking dynamically modulates visual sensitivity by contributing two new findings: first, that walking speed modulates perceived synchrony of audio-visual stimuli, and second, that modulations within the step-cycle extend to multisensory synchrony judgements which peak in the swing phase of each step.

## Introduction

Movement characterises our experience of the environment, as we shift our gaze, adjust our posture, and move about the world to achieve our behavioural goals (Noë, 2004). Despite this interplay between the sensory and motor systems, the vast majority of past research within experimental psychology and perception has focused on highly restrained experimental settings. These settings have been critical for building our understanding of human perception, but they cannot provide a complete account of how our senses function during dynamic real-world behaviour. The present study addresses this deficit by examining a critical perceptual challenge – that of judging multisensory synchrony – while engaged in one of the most common natural behaviours: locomotion. Specifically, we compare audio-visual synchrony perception while walking at a natural walk speed or at a slower speed, and we test for systematic changes in synchrony perception occurring at different phases within the step cycle.

Audio-visual events are commonly encountered in natural behaviour and careful evaluation of the relative timing of each modality’s response is essential in a multisensory brain. It is needed to determine whether two sensory signals arriving at a similar time are linked to a common external cause and should be integrated, thereby speeding reaction times and improving precision (Ernst & Bülthoff, 2004). This has clear functional benefits and can be critical for survival. For example, integrating the sound and location of an approaching car helps optimise the safe crossing of a busy road; and weak unimodal signals that may have gone unnoticed become salient when integrated. Equally importantly, however, is signal segregation. If two signals are sufficiently offset in time, it suggests they arise from distinct external causes and thus they should be kept segregated to avoid spurious binding of unrelated signals that could lead to maladaptive behavioural responses.

Accurately determining the relative timing of external auditory and visual events is not a trivial problem. There is a series of temporal factors, both external and internal, that differentially affect each modality (Alais et al., 2010). First, the travel time of the physical stimulus from source to peripheral receptors is very sluggish in the case of sound (∼3 ms per metre) but it is near-instantaneous for light. Second, once a signal arrives at the receptors, transduction times differ for vision and sound. In audition, the physical-to-neural transduction is very rapid (1-2 ms: (Corey & Hudspeth, 1979; King & Palmer, 1985)) whereas visual transduction is much slower by a factor of ∼20x (Lamb & Pugh, 1992; Rodieck, 1998). In the case of vision, latencies are further modulated by adaptation, intensity and duration (Lennie, 1981). As these asymmetries are not fixed, they require that temporal integration windows for perceived simultaneity remain flexible, for example by accommodating for auditory travel time in the case of distant audio-visual events (Alais & Carlile, 2005; Kopinska & Harris, 2004; Sugita & Suzuki, 2003).

Another factor affecting the perceived simultaneity of audio-visual stimuli is attention. Previous work has demonstrated that simultaneity perception can be flexibly biassed by attention, such as when spatially attending to a visual event leads to a speeded response to the attended signal (Kanai et al., 2007; Spence et al., 2001; Zampini et al., 2005). Thus, a given perceptual or behavioural task could alter perceived synchrony if it requires focusing on one component of the multisensory pair. Other factors affecting audio-visual synchrony perception include the content of the signals, binding windows differ when using simple tones/flashes compared to audio-visual speech (Eg & Behne, 2015). The richness of the stimulus environment also raises the difficulty of synchrony perception by increasing the number of potential distractors and raising the likelihood of misbinding (Cary et al., 2024).

In the current paper, we examine whether audio-visual synchrony perception might also be influenced by another factor: the simple act of walking. Recent work has used time-resolved perceptual probes to demonstrate that visual sensory processing undergoes continual dynamic change during locomotion (Davidson et al., 2024). More specifically, visual detection performance was shown to oscillate in a rhythm entrained to the phases of an individual’s stride cycle. During the swing phase of the stride, there was an increase in sensitivity and faster reaction times to visual stimulus probes. During the stance phase, the pattern was reversed with poorer sensitivity and slower response times. Here, we extend this approach to investigate audio-visual synchrony perception. Specifically, we test whether synchrony perception varies when comparing natural and slow walking speeds, and whether synchrony modulates within the dynamic phases of each step. Recent work has shown that the speed of self-movement can interact with timing judgements (De Kock et al., 2021; Wiener et al., 2019). Consequently we also predict that the propulsive actions involved with each step will modulate synchrony judgements as actions executed close in time to an audio-visual stimulus affect both subjective synchrony and the width of the synchrony window (Arikan et al., 2017; Benedetto et al., 2018).

To preview our results, we find that walking slowly increases overall synchrony judgements and the width of the synchrony window, demonstrating a reduction in the precision of audio-visual perception compared to when walking at a natural pace. Synchrony perception also decreased around the stance phase of the step cycle, with a greater likelihood of perceiving synchrony during the swing phase of each step.

## Methods

### Participants

We recruited 23 healthy volunteers via convenience sampling from The University of Sydney’s undergraduate psychology cohort. All participants had normal or corrected to normal vision, and provided informed consent before receiving either 20 AUD or course credit for their time. All research procedures were approved by the University of Sydney Human Research Ethics Committee (HREC 2021/048). Two participants were excluded for failure to understand the task, resulting in a final sample of 21 volunteers (13 female, Mean age 23.0 years, SD= 4.9).

### Apparatus and virtual environment

We built a virtual environment in Unity (version 2020.3.14f1), with the SteamVR plugin (ver 2.7.3, SDK 1.14.15), displayed using a DELL XPS 8950, with a 12th Gen INtel Core i7-12700K 3.60 GHz processor, running Microsoft Windows 11. The environment was similar to our previous studies investigating the effects of walking on visual perception (Davidson et al., 2023, 2024), and consisted of an open woodland, through which participants walked a 9.5 m path at two predetermined walking speeds. Participants walked the 9.5 metre path at ∼0.65 m/s in the slow condition or ∼1.1 m/s in the natural speed condition. Both speeds were chosen after extensive pilot testing, to be consistent with our prior work and previous research estimating preferred natural walking speeds over unfamiliar terrain (Davidson et al., 2024; Hausdorff et al., 1996; Matthis et al., 2018). Participants were wearing the HTC Vive Pro Eye head mounted display (HMD), which contains two 1440 x 1600-pixel (3.5” diagonal) AMOLED screens (110 degree FOV, refresh rate 90 Hz). Participants carried two wireless hand-held controllers to submit responses and self-pace trial progression, as well as the wireless adapter kit (130 gram weight) and a lightweight shoulder bag with a battery that powered the wireless device. Five HTC base stations (v 2.0) were used to record the three-dimensional position of the HMD while walking, which was used in offline analysis to determine step-cycle phase.

### Procedure

After providing informed consent at the start of the experiment, each participant was familiarised with the VR apparatus and virtual environment before beginning a first practice block which contained 5 stationary trials, as well as a sequence of easy trials (alternating easy “same” and “different” types, at 40 and 400 ms SOA respectively) to familiarise the participants with the response mapping. On all trials, participants were instructed to respond “same” or “different”, via right or left click of the index finger on the wireless controllers. On all trials, audio-visual stimulus pairs were presented at one of seven fixed stimulus-onset-asynchronies (SOAs). These were −320, −140, −50, +40, +130, +220, +400 milliseconds, with negative representing auditory leading stimuli, positive for visual leading stimuli. These values were selected after pilot testing to finely sample the centre of the synchrony function, with easy trials at the extrema to enable reasonable Gaussian fits to the data. To arrive at these SOAs, we began with a set defined as [±360 ±180 ±90 0] ms and then added an offset of 40 ms. The reason for this was to align the centre of the SOA set closer to the true point of subjective synchrony, which typically occurs when vision has a lead of about 30-40 ms relative to audition (Arrighi et al., 2005; Galton, 1885, 1899; Welford, 1980) owing to the longer transduction and latency of vision and it has been shown that auditory signals activate midbrain neurons faster than visual signals by about 50 ms (King & Palmer, 1985). Thus, setting the middle value of the set of SOAs to 40 ms provides a good estimate of the true point of subjective synchrony.

Each experimental block contained 20 trials, within which all trials were the same walking pace (either slow or natural speed). There were 3 slow walking and 6 natural walking blocks, and the entire experiment was completed in ∼1 hour. Each trial required the participants to walk the 9.5 m path over 9 seconds in the normal speed condition (velocity = 1.06m/s ≈ 1 m/s), or 15 seconds in the slow speed condition (velocity = 0.63m/s ≈ 0.6m/s, approximately 60% of the speed of the normal condition), while performing the task.

### Stimuli

On all trials participants were positioned behind a walking guide (10 cm x 10 cm cube) at approximate waist height, which traversed the length of the virtual path at either the slow or natural walk velocity. The walking guide served to ensure participants walked at a near constant velocity during walking trials, and determined the location of the visual stimulus on all trials. The visual stimulus was a large (5 cm radius) disc located 10 cm above the walking guide, which changed colour - flashing white (RGB: 1,1,1) for 20 ms. Auditory stimuli were bursts of white noise 20 ms duration, ramped over 2 ms at onset and offset. In each walking trial, Participants were presented with a maximum of 6 audio-visual stimulus pairs in the normal speed condition, and maximum of 10 pairs in the slow walking condition, such that the time allowed to respond between presentations was constant across conditions. We wish to note that both stimuli were easily perceivable, suprathreshold flashes and tones, and our main dependent variable is the perceived synchrony of these audio-visual stimulus pairs.

### Verification of SOA’s

To verify the onset of auditory and visual stimuli, the output of a photodiode attached to the HMD screen was compared to the analogue output driving the in-built HTC-Vive Pro Eye headphones, recorded concurrently using analogue inputs on a Focusright Scarlett Audio Interface. The voltage time-series were then compared using Audacity to confirm stimulus-onset timing and adjust for consistent delays. Before data collection, a significant but stable delay of 77 ms ±12 ms was found on the audio stream relative to the visual stimuli, and so a latency correction was applied in the Unity scripting to account for this, after which the empirically observed SOA’s were found to be reliably correct to approximately ±11 ms (where 11 ms is the frame duration of the 90 Hz visual display).

### Gait extraction from head position data

To test whether the phase of an individual’s step-cycle influenced synchrony judgements, we applied a peak detection algorithm to the time-series of head position data. We first detected troughs in the vertical head position data which correspond to the double support stance phase when both feet are on the ground (Gard et al., 2004; Hirasaki et al., 1999; Moore et al., 2001; Pozzo et al., 1990). We next epoched individual steps based on the trough-to-trough time-points and normalised step-lengths by resampling the time-series data to 200 data points (0-100% stride-cycle completion, incrementing in steps of 0.5%).

Stimulus onsets were then allocated into their respective step-cycle quintiles, corresponding to whether the first stimulus in an audio-visual pair was shown in the 1-20%, 21-40%, 41-60%, 61-80%, 81-100% resampled percentile bins. To increase the trial counts per quintile trials were aggregated from both walking speeds.

### Model fitting

To assess the sensitivity of synchrony judgements while walking, we applied a two-step procedure. First, the proportion “same” responses for each SOA were quantified per participant per walking speed, before a Gaussian model was fit to the individual level data. We fit using the standard Gaussian function:

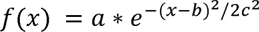

Where a is a constant, e is Euler’s number, b is the SOA at peak of the synchrony distribution and c is distribution’s standard deviation. For each participant, we fit a Gaussian to their average per SOA using a non-linear least squares method (max 2000 iterations), and retained the peak centre (b) and standard deviation (c) for statistical comparisons between walking speeds. To improve fit strength and focus on our parameters of interest (mean and standard deviation), we normalised the height of each participant’s synchrony function to 1, by dividing by their maximum proportion of “same” responses.

We additionally performed a series of asymmetric Gaussian fits to quantify whether the auditory leading or visual leading stimulus pairs interacted with walking speed. These used the same equation, but separately quantified the standard deviation for the left-hand side relative to the Gaussian mean (LHS, auditory leading), or right-hand side relative to the Gaussian mean (RHS, visual leading) of each participant’s data.

### Mixed-effects analyses

We investigated the effect of step-cycle phase on synchrony judgements, focussing on changes to the proportion of ‘same’ responses at the shortest SOAs (−50ms, +40ms). We focussed on these short SOAs to ensure that the entire stimulus sequence would be approximately 60-70 ms in duration (e.g. 20 ms flash, 20 ms gap, 20 ms tone for the 40 ms SOA, with 60 ms total duration), thus enabling an analysis of how relatively brief step-cycle phases would influence performance.

Owing to the few number of targets at this subset of SOAs we combined both walk speeds to increase the statistical power of our analysis (Combined speeds, total trials at SOA −50 ms *M =* 192.04, SD = 10.95; SOA +40ms, *M* = 195.90, SD = 13.97). Assigning these short SOA trials into step-cycle quintiles resulted in approximately 36 trials per step-cycle quintile (SOA −40 ms Mean across quintiles = 36.11, *SD* = 1.07; SOA +50 ms, M = 36.69, *SD* =1.37).

We observed quadratic trends in the proportion of ‘same’ responses at these short SOAs depending on step-cycle quintile, and formally tested for quadratic effects using stepwise mixed-effects analyses. This analysis compared whether models including only a linear effect (P(same) ∼ step quintile + (1|Participant)) or additional quadratic fixed effect (P(same) ∼ step quintile^2^ + step quintile + (1|Participant)) were better than a simple model including only random effects (intercepts) per participant (P(same) ∼ 1 + (1|Participant)). We compared the goodness-of-fit of each model using likelihood ratio tests, and report when the quadratic or linear model was a significantly better fit than the basic model, including coefficients for fixed effects (β), their 95% Confidence Intervals (CIs).

## Results

We quantified audio-visual synchrony judgements during walking at slow and natural speeds to investigate the impact of everyday actions on multisensory perception. **Figure 1** displays a representation of the basic virtual environment and task design.

**Figure 1-.**
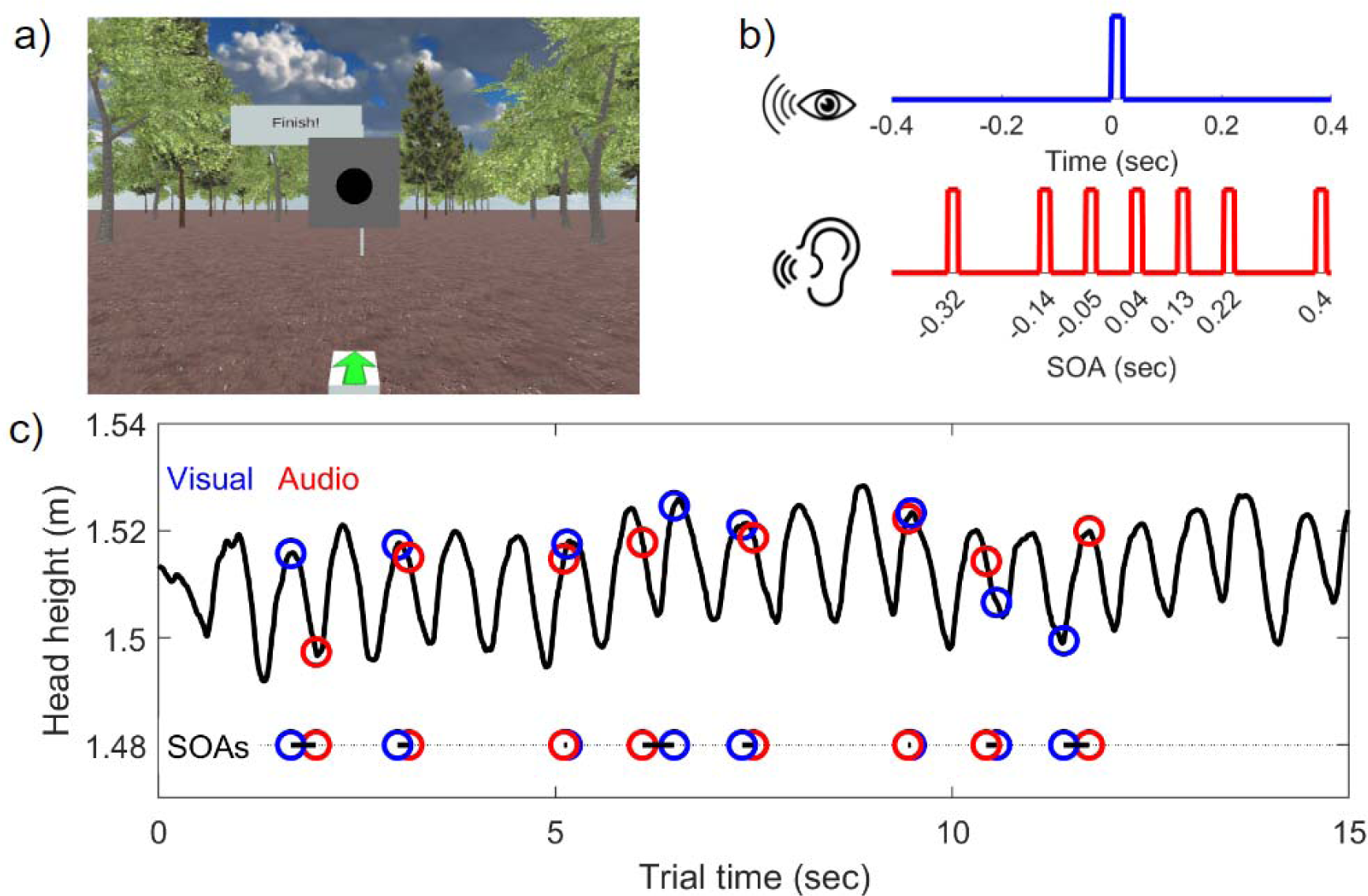
Audio-visual synchrony judgements while walking at natural and slow speeds. **a)** First-person view of the 3D environment participants walked through while providing synchrony judgements. **b)** Example stimulus timing showing the stimulus onset asynchronies used in the present experiment, ranging from −0.32 seconds (audio leading visual), to 0.4 seconds (visual leading audio). **c)** Example change in head position recorded during a single trial at slow walking speed. Vertical head height displays a roughly sinusoidal pattern where the peaks and troughs correspond to the approximate swing and stance phases of the stride-cycle (solid black line). Visual onsets are shown with blue markers, auditory onsets are shown with red markers, overlaid on the head position to display their occurrence in various phases of each step. The SOA is displayed on the horizontal line at bottom.

### Slow walking reduces the sensitivity of auditory-visual synchrony judgements

In studies of multisensory synchrony, plotting the proportion of synchronous percepts per SOA typically results in a Gaussian bell-shaped function (Vroomen & Keetels, 2010). The mean (or peak) of the function represents the point of subjective simultaneity (PSS), and the width of the function – particularly the slope of the rise and fall – provides a measure of the sensitivity of synchrony judgements. A very narrow function represents a tight SOA range at which two stimuli will be perceived as simultaneous, and thus higher sensitivity. Conversely, a wider function represents a broader range at which two sequential stimuli are perceived to be simultaneous, and reduced sensitivity.

The average proportion of synchronous percepts when walking at a slow and natural pace are displayed in **Figure 2A**. On visual inspection, a wider function is observed for the slow walking condition which we formally quantified by comparing the standard deviation of the Gaussian fits obtained per participant at both walking speeds. Synchrony functions were significantly wider when walking slowly, compared to when walking at a natural speed (*t*(20) = 3.88, *p* < .001, *d* = .85). There was no significant difference between the mean (PSS) of the synchrony functions at both walking speeds (*t*(20) = .4, *p* = .7, *d* = .09). We also conducted a 2 x 7 repeated-measures ANOVA to assess the overall proportion of synchronous percepts in the natural and slow walk speeds over the seven SOAs. This revealed a significant main effect of walk speed (*F*(1,20) = 11.674, *p* =.003, η_p_^2^ = 0.369), with slow walk speed producing a higher proportion of “same” responses. As is clear from **Figure 2A**, the main effect of SOA was also significant (*F*(6,120) = 110.643, *p* <.001, η_p_^2^ = 0.847). The walk speed x SOA interaction was not significant.

**Figure 2.**
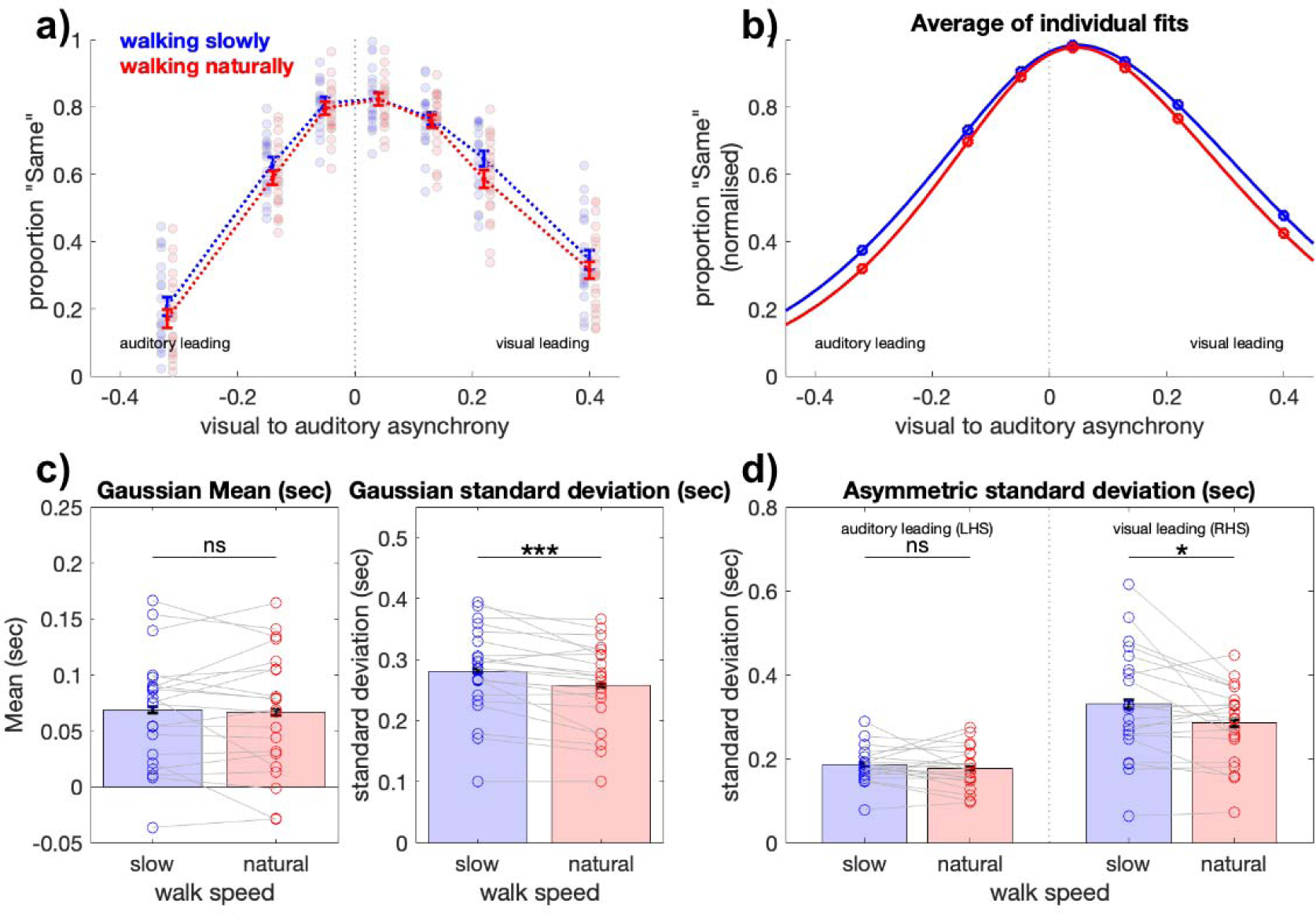
Walking at slow speed decreases the sensitivity of synchrony judgements. **a)** Group average proportion ‘same’ responses per SOA. Blue and red colours represent slow and natural walking speeds respectively. Error bars represent ± 1 SEM, corrected for within participant comparisons (Cousineau, 2005). **b)** Average of participant-level Gaussian fits at slow and natural walking speeds. Each fit was performed after normalising to the maximum proportion ‘same’ per participant, to improve the fit strength for the Gaussian mean and standard deviation parameters of interest. **c) Left:** Gaussian mean per participant was not significantly different at slow and natural walking speeds. **Right:** Gaussian width per participant significantly reduced when walking at natural speeds. **d)** Asymmetric Gaussian fits (fitting the LHS and RHS separately) show the visual leading stimuli on the RHS were significantly affected by walking speed.*** p< .001, * p < .05 Bonferonni corrected

We additionally calculated the standard deviation of Gaussians fit to either the left hand side (LHS) or right hand side (RHS) of the Gaussian mean at both walking speeds. This analysis serves to quantify any asymmetry in the Gaussian profile, which represents an interaction between walking speeds and the type of leading stimulus (auditory leading or visual leading stimulus pairs on the left and right, respectively). These tests of asymmetric fits revealed that on average, the synchrony functions were significantly wider to the right of the Gaussian mean (i.e., vision leading stimuli: RHS standard deviation *M=* 0.30, *SD* = .09) compared to the left of the Gaussian mean (i.e., auditory leading stimuli: LHS *M* = 0.18, *SD =* .04; *t*(20) = −6.76, p<.001, *d* = −1.48). A 2×2 repeated measures ANOVA (slow/natural speed x auditory leading/visual leading standard deviation) indicated significant main effects for the left/right asymmetry (*F*(1,20) = 39.75, *p* <.001, η_p_^2^ = 0.67), and walking speed (*F*(1,20) = 12.59, *p* = .002, η_p_^2^ = .39), with no significant interaction (*F*(1,20) = 2.17, *p* = .16). Post-hoc tests revealed the main effect of walking speed was driven by a change to the width on the RHS of the Gaussian (*t*(20) = 3.10, *p* = .02, *d* =0.2; Bonferroni corrected for comparing a family of 6).

In summary, audio-visual synchrony judgements are biassed by whether the visual or auditory stimulus was leading in the pair, with visual leading stimuli perceived with poorer sensitivity. All stimulus pairs were impaired when walking slowly compared to when walking at a natural speed, an effect that was driven by an increase in the width of the synchrony function at slow walking speeds for visual-leading stimuli.

### Slow walking increases reaction times

We next analysed whether walking speed and the modality of the leading stimulus also affected reaction times. While overall performance and reaction times often correlate in speeded tasks such as ours, they can offer unique and complementary information capturing distinct aspects of cognitive processes (De Boeck & Jeon, 2019). For example, a change in sensitivity without a change in reaction time could imply an objective increase in response error with no change in the perceived difficulty of the task (such as the misattribution of button-presses). Alternatively, a change in sensitivity and increase in reaction times could imply that walking slowly altered the perceived difficulty of the task, inducing a reaction time delay.

**Figure 3A** displays the overall reaction time from the first stimulus per SOA when walking at slow and natural speeds. Like the proportion ‘same’ responses, reaction times displayed a roughly symmetrical Gaussian shape, although interestingly, reaction times were fastest overall for the stimuli with shortest SOAs. This pattern was despite the large suprathreshold asynchrony presented at maximum SOAs, and our instructions to participants that were to respond as quickly as possible. Responses could be made before the onset of the second stimulus during these larger SOAs, yet clearly that was not the dominant strategy. One explanation for this result is that participants may have waited for both stimuli prior to committing to a response. If this were the case, then large SOAs may have also primed a rapid response when measured relative to the onset of the second stimulus. **Figure 3C** displays the reaction times from the second stimulus, (i.e. subtracting the absolute SOA from the reaction times in **Figure 3A**), from which several interesting features emerge.

**Figure 3.**
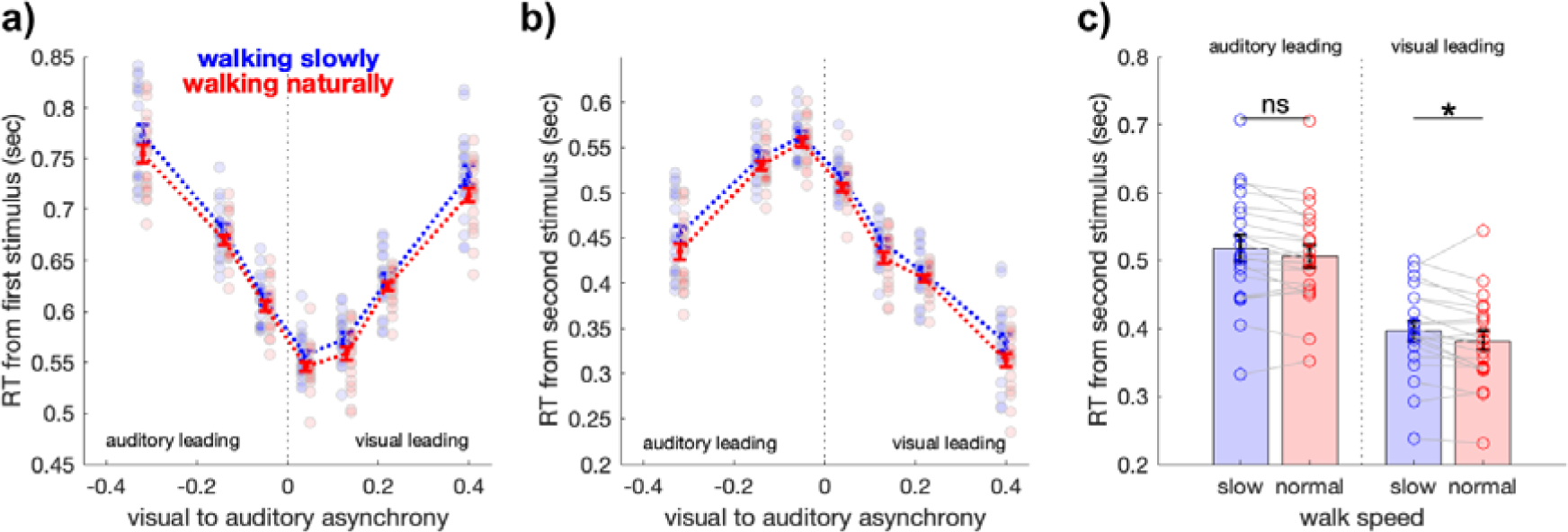
Walking slowly increases reaction times. a) Group average reaction times measured from the onset of the first stimulus. Blue and red colours represent slow and natural walking speeds respectively. Error bars represent ± 1 SEM, corrected for within participant comparisons (Cousineau, 2005). b) Group average reaction times when measured from the onset of the second stimulus. c) Average reaction times in b, when walking slowly (blue) or naturally (red), when the stimulus pair was auditory leading (SOAs −320, −140, −50 ms) or visual leading (SOAs +40, +130, +220, +400 milliseconds). Error bars represent ± 1 SEM, participant data shown with circles, and connected per participant with grey lines. * p < .05 Bonferonni corrected

First, reaction times were fastest overall for stimuli with larger SOAs, implying a priming of rapid responses in those trial types. Second, the shortest SOAs, for which the maximum proportion of ‘same’ responses were observed, resulted in longer reaction-times. There was also a clear asymmetry in reaction-time behaviour for auditory leading compared to visual leading stimulus pairs, despite the symmetrical pattern of response-times when measured relative to the first stimulus (cf. **Figure 3A**). Large differences in reaction times were also clear between visual leading and auditory leading stimulus pairs, despite objectively similar absolute SOAs. For example, for auditory leading SOAs of −0.14 sec, reaction times (*M* = 0.53, *SD*=.08) were over 100 ms slower than visual leading SOAs of +0.13 sec (*M* =0.42, *SD =* .07), despite an objective difference in absolute SOA of only 10 milliseconds (*t*(20) = 11.82, *p* < .001, *d* = 2.58).

We proceeded by quantifying the effects of walking speed on these reaction times with a 2×2 repeated measures ANOVA (slow/natural speed x visual/auditory lead). There was a significant main effect for the modality of the leading stimulus (*F*(1,20)= 322.19, *p* < .001, η_p_^2^ = 0.94), and walking speed (F(1,20) = 9.71, p = .005, η_p_^2^ = .33), with no significant interaction (*F*(1,20) = 0.12, *p* = .74). Post-hoc tests revealed the main effect of walking speed was significant only for the RHS visual leading stimuli (*t*(20) = 2.87, *p*= .04, *d*= .59; LHS, *p* = .1; Bonferroni corrected for comparing a family of 6).

Overall, these results indicate that slower reaction times accompany the decrease in sensitivity that occurs when walking slowly (cf. **Figure 2**). When measured from the first stimulus in a pair, reaction times were fastest when SOAs were minimal (**Figure 3A**). On average, however, participants withheld their responses until after both stimuli had been presented, resulting in fastest reaction times for larger SOAs when measuring reaction times from the second stimulus in a pair (**Figure 3B**). There was also a significant advantage for visual leading stimulus pairs, which were responded to faster than auditory leading alternatives. However, responses to visual leading stimuli were significantly slower when walking slowly.

### Step-cycle phase modulates synchrony judgements at shortest SOAs

Recent evidence has demonstrated that visual detection performance is rhythmically altered throughout the step-cycle. In this previous work, changes in head position while walking in wireless VR were used to define the approximate swing and stance phases of locomotion. Here, we sought to extend this line of work to investigate whether step-cycle phase would also impact upon judgements of audio-visual synchrony. For this analysis, we divided each participant’s step-cycle into five quintiles (binning according to the 1-20%,21-40%, 41-60%, 61-80% and >80% step-cycle percentiles) and focused on the shortest SOA condition (+40 ms) which was closest to the point of subjective simultaneity (PSS), and ensured that the entire stimulus sequence would be 60 ms in duration (e.g. 20 ms visual flash, 20 ms gap, 20 ms auditory click = 40 ms SOA, 60 ms total duration). This focus ensured that both stimuli would be shown in either a single step-cycle quintile or neighbouring step-cycle quintiles (see Methods), facilitating an analysis of how step-cycle phasemay impact perceived simultaneity judgements. We note that analysing the impact of step-cycle phase for longer SOAs is also possible but problematic to interpret, as the asynchronous stimuli will be presented in separate phases or steps as SOAs increase. For comparison, we present changes in simultaneity judgements across the three shortest SOAs (−50, +40, +130; **Figure 4**).

**Figure 4.**
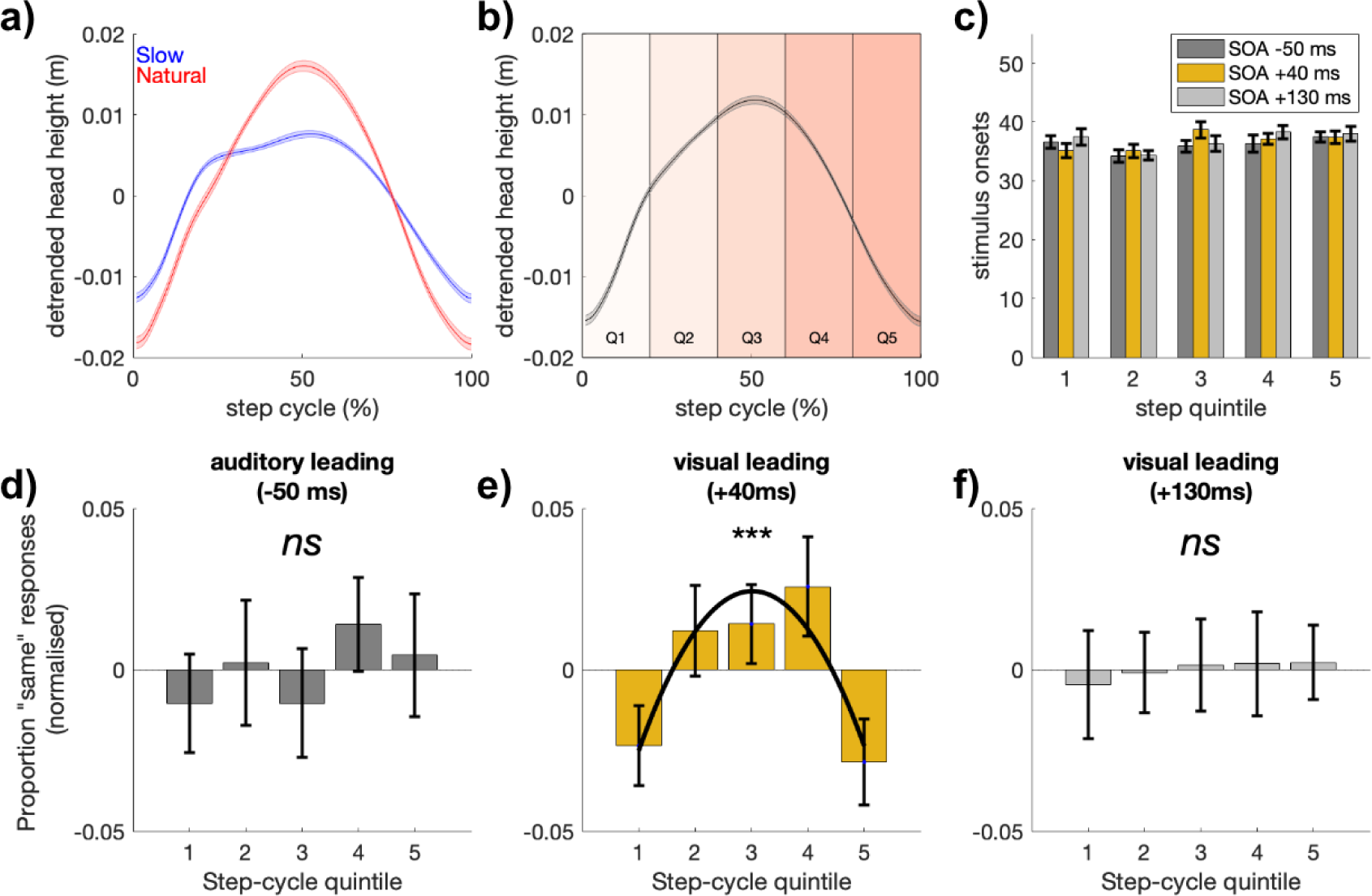
Proportion ‘same’ responses at short SOAs are modulated by step-cycle quintile. a) Grand average detrended change in head-height at slow (blue) and natural (red) walking speeds during a single step. Note to facilitate step-cycle comparisons, each step is normalised to 1-100% step cycle completion. b) Average head-height using both speeds over a single step, with step-cycle quintiles overlayed. c) Average trial count per step-cycle quintile for the three shortest SOAs (−50, +40, +130 ms, shown in dark-grey, yellow, light-grey, respectively). d-f) Proportion same responses per this selection of SOAs. Only the shortest SOAs closest to the PSS (+40 ms) were significantly modulated by step-cycle quintile.

**Figure 4** displays the proportion of ‘same’ responses for these shortest SOAs, including SOA +40 ms which was closest to the point of subjective simultaneity. At this SOA the proportion of ‘same’ responses decreased around the time of footfall. We formally tested for this quadratic tend using mixed-effects models and likelihood ratio tests, and found a quadratic model (with random intercepts per participant) was a significantly better fit than the basic model that included only random intercepts per participant (quadratic vs basic: ^2^(2) = 11.11, *p* = .004, β = −0.012 [-0.02, −0.005]) as well as a simple linear model (quadratic vs linear: χ (1) = 11.10, *p* = 8.6×10). The two neighbouring SOAs (auditory leading −50 ms, visual leading +130 ms) demonstrated no impact of step-cycle quintile on synchrony judgements, as neither the linear or quadratic models were a better fit than the basic model (*p*s > .4).

In summary, at the shortest SOAs closest to the point of subjective simultaneity, synchrony judgements are altered by step-cycle phase in a quadratic manner. Judgements of synchrony for SOAs at the point of subjective simultaneity decrease around the time of footfall.

## Discussion

Here we investigated perception of audio-visual synchrony during locomotion, testing whether synchrony perception would be affected by walking speed and whether it would vary with the phase of the stride cycle. Participants walked at slow and natural walking speeds in a wireless virtual-reality environment indicating whether pairs of audio-visual stimuli were synchronous or not, responding as quickly as possible. We compared the proportion of ‘synchronous’ judgements and reaction-times to stimuli presented at randomly jittered intervals as participants walked a straight and level path. Overall, the effect of slow walking was to increase the probability of ‘synchronous’ judgements and to increase reaction time. An asymmetry was also observed such that walking slowly most strongly affected the visual-leading rather than auditory-leading stimulus pairs. To examine whether synchrony perception modulated with the phases of the step cycle, we analysed the shortest SOA which corresponded closely to the point of subjective simultaneity and found that the proportion of synchronous responses rose and fell across the course of a single step. Step cycle phase modulated responses in a quadratic manner, with synchronous responses being lower around the time of footfall at the start and end of each step and higher in the swing phase. Together these results reveal that walking speed and step cycle phase both affect the perception of multisensory synchrony and extend recent unisensory work showing that visual detection is modulated by the stride cycle while walking at a natural speed (Davidson et al., 2024).

### Walking at slow speed impairs audio-visual synchrony judgements

A key finding was that synchrony perception depended on walking speed, with the width of Gaussian functions fit to the proportion of synchronous responses being broader when walking slowly. The overall proportion of synchronous percepts was also higher, and reaction times slower during the slow walk condition. This broadening of the synchrony function in slow walking is effectively a decrease in sensory precision arising from a degraded ability to distinguish the offset of the visual and auditory events.

What might underlie this change in sensory precision during walking? Relative timing judgements have previously been explained by independent component models (Sternberg & Knoll, 1973). In these models, a change in synchrony width would be modelled by shallower psychometric functions for the auditory and visual components respectively (see **Figure 5**). In our analysis, there was no difference in the point of subjective simultaneity (i.e. the Gaussian mean) between slow and natural walk speeds, meaning the effect of slow walking was confined to a change in the slope of the component psychometric functions, rather than a shift in their means.

**Figure 5.**
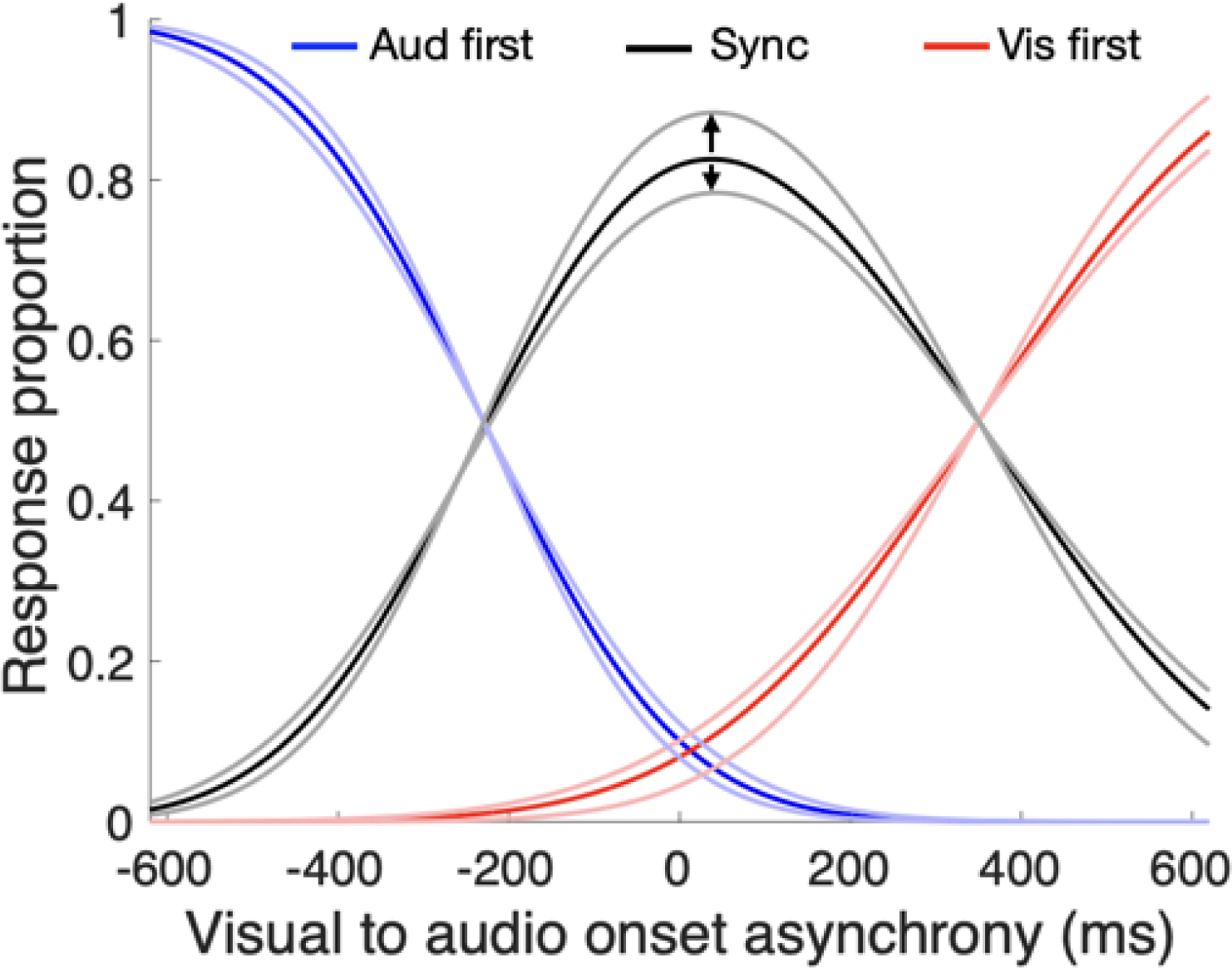
A modulation of proportion synchronous responses driven by changing auditory and visual timing precision over the step cycle. Based on independent channels models (e.g., (Sternberg & Knoll, 1973)) commonly used to explain relative timing, variations in the slope of component psychometric functions will alter the peak synchrony value. If component slopes were steepest during the swing phase of the step cycle, as implied by previous results (Davidson et al., 2024), then peak synchrony judgements would also modulate with the step cycle and peak in the swing phase, as observed in Figure 4E.

The main factors that cause a broadening of psychometric functions, and thus degraded precision, are a poorer signal-to-noise ratio in the stimulus or a reduction in attentional resources applied to the task. The act of locomotion likely creates a degree of noise and disruption to perception, and this would apply similarly to slow and natural walking. The explanation of the broader synchrony function during slow walking may have to do with increased attentional demands and effort required to walk at a slower than natural pace.

Indeed, humans exhibit a remarkably narrow distribution of step rate when walking at a natural speed that is centred on a mean value close to two steps per second, corresponding to a peak in biomechanical efficiency (McNeill Alexander, 2002; Zarrugh et al., 1974). Walking faster than this preferred rate requires more effort and, on a naïve view, slow walking would seem to require less effort than the preferred rate. Curiously, however, this is not the case. There is a U-shaped function of effort over walk speed, with both slow and faster speeds requiring more effort (Ralston, 1958) When walking slowly, the attraction to this inherently efficient and preferred speed must be overcome by a wilful, top-down inhibition of the low-level, spinal circuits that drive the walking oscillation (Schubert et al., 1999). We speculate that this effort draws away cognitive resources that would otherwise be free to be deployed on the synchrony judgement. The effect of deploying less cognitive resources could reduce sensitivity on the task either by broadening the component psychometric functions (as shown in **Figure 5**) or by reducing signal salience. For example, participants have shown a greater propensity to judge stimuli as synchronous at low intensity than at high intensity, thus affecting the width of the synchrony function (Krueger Fister et al., 2016). Consistent with these accounts, reaction times were also slower in the more effortful condition of slow walking. Future experiments could arbitrate between the attentional and effort accounts by measuring synchrony perception during locomotion employing a 2×2 factorial design, comparing low/high task demands with low/high efforts of locomotion.

### Broad synchrony functions during action

It is notable that our estimates for audio-visual synchrony derived from Gaussian fits are broader than is typically reported. In this study, we fitted an asymmetric Gaussian to the normalised mean data per participant (i.e., separate standard deviation estimates were fitted to the left and right sides of the synchrony function, see **Figure 2B**) based on findings showing that the left and right sides of the synchrony function often show a considerable asymmetry (García-Pérez & Alcalá-Quintana, 2012; Yarrow et al., 2011). We therefore defined the temporal band width of perceived synchrony as the sum of the left- and right-side Gaussian standard deviations. Using this metric, we find full-width estimates of 457 ms and 520 ms for the natural and slow walk speeds, respectively. These values are large when compared to the majority of other studies of audio-visual synchrony conducted with passive participants. A survey of passive studies shows a mean synchrony window (twice the Guassian standard deviation) of about 335 ms, although values vary with task. It is inferred to be about 200 ms by varying asynchronies in the McGurk effect (van Wassenhove et al., 2007), as high as 400 ms using the stream/bounce illusion (Shimojo & Shams, 2001), 300-400 ms using a standard flash/click stimulus (Alais et al., 2017; Lennert et al., 2021; Noel et al., 2018; Van der Burg et al., 2018), and 300 ms using realistic dynamic audio-visual events (Eg & Behne, 2015). Our values of 457 ms and 520 ms are considerably higher than other reports.

The most salient feature to explain this key point of difference is our use of active participants. This appears to be critical, as the only other study to measure the perception of audio-visual synchrony in an active context reported a similarly large value of 520 ms (Arikan et al., 2017). In that study, the effect of action upon synchrony percepts was assessed by triggering audio-visual stimuli by a button press. They found that a contiguous action broadened the window of perceived synchrony and that introducing a delay between the action and the stimuli caused the window to become narrower. Based on this observation, it appears that action around the time of a near-synchronous audio visual stimuli increases the likelihood that they will be judged as synchronous. Here we can extend this result by applying a similar interpretation to our participants, who made synchrony judgements during the continuous action of walking. Thus, the only two studies of audio-visual synchrony in active observers agree in reporting larger windows for perceived synchrony than is typical of passive studies (reviewed above, with a mean of 335 ms). It remains to be seen whether this effect on audio-visual synchrony judgements during locomotion would generalise to synchrony of other multisensory stimuli such as tactile-vision and tactile-auditory stimuli.

Why would action impair synchrony perception? Executing an action is known to produce periods of degraded perception around the time of the action. This is clear from the literature on saccades which has demonstrated strong suppression of vision around the time of saccades as well as distortions of perceived space and time (Ross et al., 2001). The phenomenon of tactile suppression is similar and well established – a loss of tactile sensitivity that occurs when reaching for objects or making other goal-directed movements. This effect is also known as ‘movement-related gating’ and its time-course suggests that it serves to suppress movement-related feedback during action, with suppression released in time for contact and grasping so that perception is optimised when needed (Chapman et al., 1987; Juravle et al., 2010). Walking also involves goal directed actions, yet much less is known about how the act of locomotion impacts on perception. One recent study used a new method to probe perception continuously over the stride cycle and found visual sensitivity modulated approximately sinusoidally, peaking in the swing phase and reaching a minimum during stance (Davidson et al., 2024). Although focused more on visual-motor control, another recent study continuously measured reaching accuracy to a visual target over the stride cycle and found similar modulations, with greater error in swing than stance phase (Davidson et al., 2023). Overall, given the many ways action interacts to constrain perception, it is not surprising that audio-visual synchrony during action would be less precise than in passive observers, even if the precise causes are not clear at this stage.

### Perceived synchrony modulates over the step-cycle

The final observation of interest was that the point of maximum synchrony modulated over the step cycle in a quadratic fashion, with peak synchrony occurring in the middle of the swing phase (**Figure 4**). Based on the independent channels models commonly used to explain relative timing (Sternberg & Knoll, 1973), variations in the slope of the component psychometric functions will alter the peak synchrony value, as illustrated in **Figure 5**. Given the prior evidence of visual performance modulating within the step cycle (Davidson et al., 2024)), and data forthcoming from our lab showing similar oscillations in auditory sensitivity (Davidson et al., *in prep*), we propose that a steepening of component slopes during the swing phase is what underlies the peak of perceived synchrony in the swing phase, as observed in **Figure 4E**.

A final point to note is that more recent versions of the independent-channels model (Sternberg & Knoll, 1973) have revised it to include a fourth parameter so that separate estimates can be obtained of the width of the synchrony function on the left and right side of the mean (García-Pérez & Alcalá-Quintana, 2012; Yarrow et al., 2011). For this reason, we fit an asymmetrical Gaussian function with separate left and right side standard deviations (see **Figure 2d**). The results revealed a significantly larger standard deviation on the right side in the slow walk condition, which ties the reduced sensitivity to the right side that is governed by visual timing precision. From these data we can conclude that there is a loss in sensitivity to synchrony perception in general during slow walking relative to natural walk speed, and, more specifically, that this loss is due to impaired sensitivity to visual timing when walking slowly.

### Conclusions

Overall these data extend recent efforts to move from the seated tradition of laboratory experiments into more ecologically plausible dynamic environments. By testing audio-visual synchrony perception during a frequent everyday behaviour, we have revealed that recent modulations in unisensory vision within the stride-cycle also extend to the perception of multi-sensory timing.

